# The transcription factor NF-YA10 determines the area explored by *Arabidopsis thaliana* roots through direct regulation of *LAZY* and *TAC* genes

**DOI:** 10.1101/2023.04.26.538431

**Authors:** Andana Barrios, Nicolas Gaggion, Natanael Mansilla, Leandro Lucero, Thomas Blein, Céline Sorin, Enzo Ferrante, Martin Crespi, Federico Ariel

## Abstract

Root developmental plasticity relies on transcriptional reprogramming, which largely depends on the activity of transcription factors (TFs). NF-YA2 and NF-YA10 (Nuclear Factor A2 and A10) are down-regulated by the specific miRNA isoform miR169defg, in contrast to miR169a. Here, we analyzed the role of the *Arabidopsis thaliana* TF NF-YA10 in the regulation of lateral root development. Plants expressing a version of *NF-YA10* resistant to miR169 cleavage showed a perturbation in the lateral root gravitropic response. By extracting novel features of root architecture using the ChronoRoot deep-learning-based phenotyping system, we uncovered a differential emergence angle of lateral roots over time when compared to Col-0. Detailed phenotyping of root growth dynamics revealed that NF-YA10 activity modulates the area explored by Arabidopsis roots. Furthermore, we found that NF-YA10 directly regulates *TAC1* and *LAZY* genes by targeting their promoter regions, genes previously linked to gravitropism. Hence, the TF NF-YA10 is a new element in the control of LR gravitropism and root system architecture.

## INTRODUCTION

Plant developmental plasticity relies on a plethora of adaptive strategies in response to the environment. The resulting root system architecture needs to ensure efficient anchor and uptake of water and nutrients. The density and length of lateral roots (LRs) expand the plant surface contact with the substrate, thus impacting the general growth of the plant. In the model species *Arabidopsis thaliana*, LR development is tightly regulated by cellular and molecular mechanisms from the very first cell division in the pericycle, through the formation of the new meristem and the emergence of the new organ from the main root. The intricate regulatory network controlling LR development includes key transcription factors (TFs) integrating internal and environmental cues (Lavenus et al., 2015). The regulatory hub formed by specific miR169 isoforms and their target TF *NF-YA2* was previously described as a modulator of root architecture. Plants resistant to the miR169-mediated down-regulation of *NF-YA2* exhibit an enhanced density of LRs explained by altered specific cell type number and greater root meristem size (Sorin *et al*., 2014). Although the related TF NF-YA10 (Leyva-Gonzalez *et al*., 2012, Zhao *et al*., 2017) is also expressed in roots and regulated by miR169, its potential role in LR development remains unexplored. In contrast, NF-YA10 was found implicated in leaf development, regulating directly IAA biosynthesis (Zhang *et al*., 2017). In addition, an enhanced stress-tolerant phenotype was described for plants overexpressing NF-YA10 and NF-YA2 (Leyva-Gonzalez *et al*., 2012). A transcriptomic analysis of both transgenic lines hinted at a functional redundancy between both NF-YA TFs with half of common deregulated genes, consistent with the high similarity exhibited by all NF-YA TFs at the protein level (Siefers et al., 2009; Petroni et al., 2012; Laloum et al., 2013). Furthermore, single homozygous mutant lines do not show a major phenotype (Zhao *et al*., 2020) except for embryo-lethality in multiple or specific single NF-YA mutants (Fornari et al., 2013; Pegnussat *et al*., 2008). The activity of the *NF-YA10* promoter revealed by the control of reporter genes indicated that *NF-YA10* is expressed in the shoot apical meristem (SAM) as well as in the main root and LRs, and is induced during phosphate deficiency, oppositely to miR169 (Leyva-Gonzalez *et al*., 2012; Sorin *et al*., 2014). Here, we challenged the hypothesis of redundancy between NF-YA10 and the closely related TF NF-YA2 by undertaking in-depth characterization of the root system architecture dynamics upon NF-YA10 deregulation. To this end, we leveraged the potential of ChronoRoot, a high-throughput automatic phenotyping system based on deep learning (Gaggion *et al*., 2021), for which we expanded here the analysis by introducing two new angle measurements as additional parameters. Plants carrying a *NF-YA10* resistant to miR169 cleavage (NF-YA10 miRres) exhibit an increased root area, due not only to a greater LR density than the wildtype (WT), but also as a result of an alteration of LR angles, suggesting a link with gravitropic responses. Furthermore, we demonstrated that NF-YA10 directly regulates *TAC1* and *LAZY* genes, previously linked to the gravitropic response of roots (Kawamoto and Morita, 2022), indicating that NF-YA10 may act as a coordinator of LR distribution in the rhizosphere, shaping the final surface of the root architecture.

## RESULTS

A phylogenetic analysis of plant NF-YA TFs indicated that this family of TFs presents four clades with internal duplications that are characteristic to the plant kingdom (**Supplemental Figure 1, Supplemental Table 1**). As previously proposed (Laloum et al., 2013), NF-YA2 and NF-YA10 (clade D) arose from a recent duplication which seems to be specific to Brassicaceae, similarly to other NF-YAs such as NF-YA1/9 (clade C) and NF-YA4/7 (clade B). In another hand, NF-YA6 and NF-YA5, NF-YA3 and NF-YA8 emerged from two duplications inside the clade A (**Figure 1A, Supplemental Table 1**). Expression studies based on transcriptional reporter *pNF-YA10:GUS* assays were used to show *NF-YA10* specific expression in the root vasculature (Leyva-Gonzalez et al., 2012; Sorin et al., 2014). In this study, we used transgenic plants expressing the translational fusion *pNF-YA10:GFP-NF-YA10mut* (miR169-resistant *NF-YA10*, called hereafter NF-YA10 miRres), to localize NF-YA10 protein expression pattern. By analyzing NF-YA10 miRres plants, we observed GFP:NF-YA10 accumulation in the nuclei of root vasculature and cortex cells, and particularly abundant at the base of the LR (**Figure 1B**) suggesting a regulatory role in the development of this organ.

**Figure 1.**
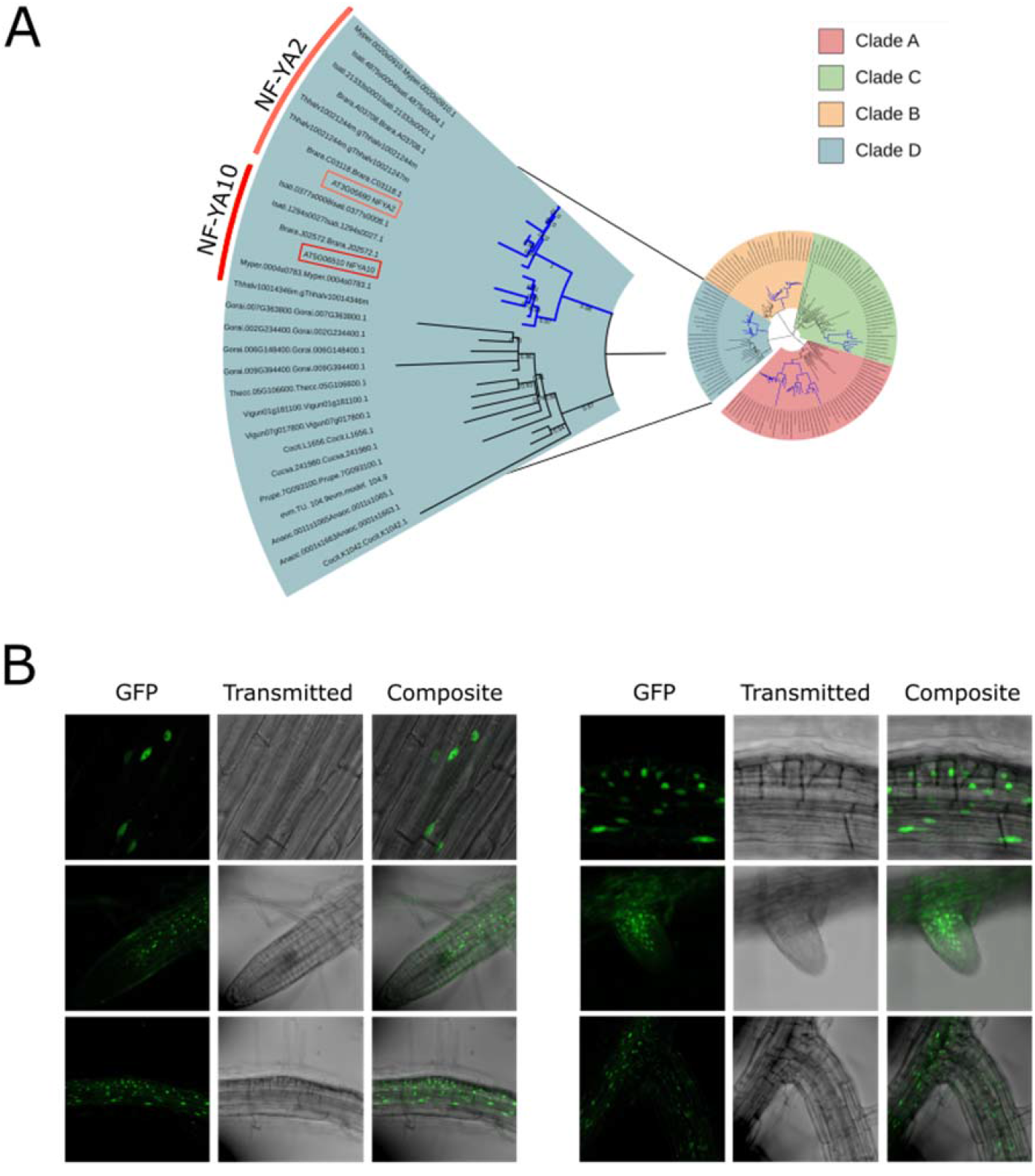
NF-YA10 diverged from a recent duplication with NF-YA2 within Brassicaceae and is expressed in nuclei of primary and lateral root vasculature cells. (A) Phylogenetic tree of NF-YAs with extended number of Malvidae and Brassicaceae, where AtNF-YA10 is represented in red and NF-YA2 in orange. The duplications in Brassicaceae are colored in blue. Only branches with bootstrap values higher than 65% are shown. (B) Localization of NF-YA10 fused to GFP in roots of pNF-YA10:GFP-NF-YAmiRres.1 8-day-old plants.

A comprehensive characterization of the root architecture dynamics was then undertaken by using ChronoRoot (Gaggion et al., 2021), i.e. comparing two independent NF-YA10 miRres lines and the WT. NF-YA10 miRres plants showed a slightly longer main root (MR) than WT, a feature observed since the germination (**Figure 2A**). Interestingly, total LR length in NF-YA10 miRres plants increased at a greater speed than the WT, showing a stronger difference than for the MR (**Figure 2B**). Therefore, although NF-YA10 miRres plants exhibit a general faster growth of the whole root system (**Figure 2C**), the relative contribution of the MR to the global root system is less significant (**Figure 2D**). Moreover, the number of LRs was significantly higher in NF-YA10 miRres plants (**Figure 2E**), resulting in an enhanced LR density (**Figure 2F**). The final root architecture of NF-YA10 miRres plants exhibited an enhanced covered surface, which is illustrated by the significantly expanded convex hull area of the full root system (**Figure 2G**). However, the density of the LRs covering the convex hull area (**Figure 2H**) did not emerge as a distinct feature between both independent lines, whereas the aspect ratio (height/width of the root system) of NF-YA10 miRres plants was significantly different from the WT, at least at day 9 (**Figure 2I**), indicating that longer LRs expand away from the MR instead of keeping closer to the MR axis.

**Figure 2.**
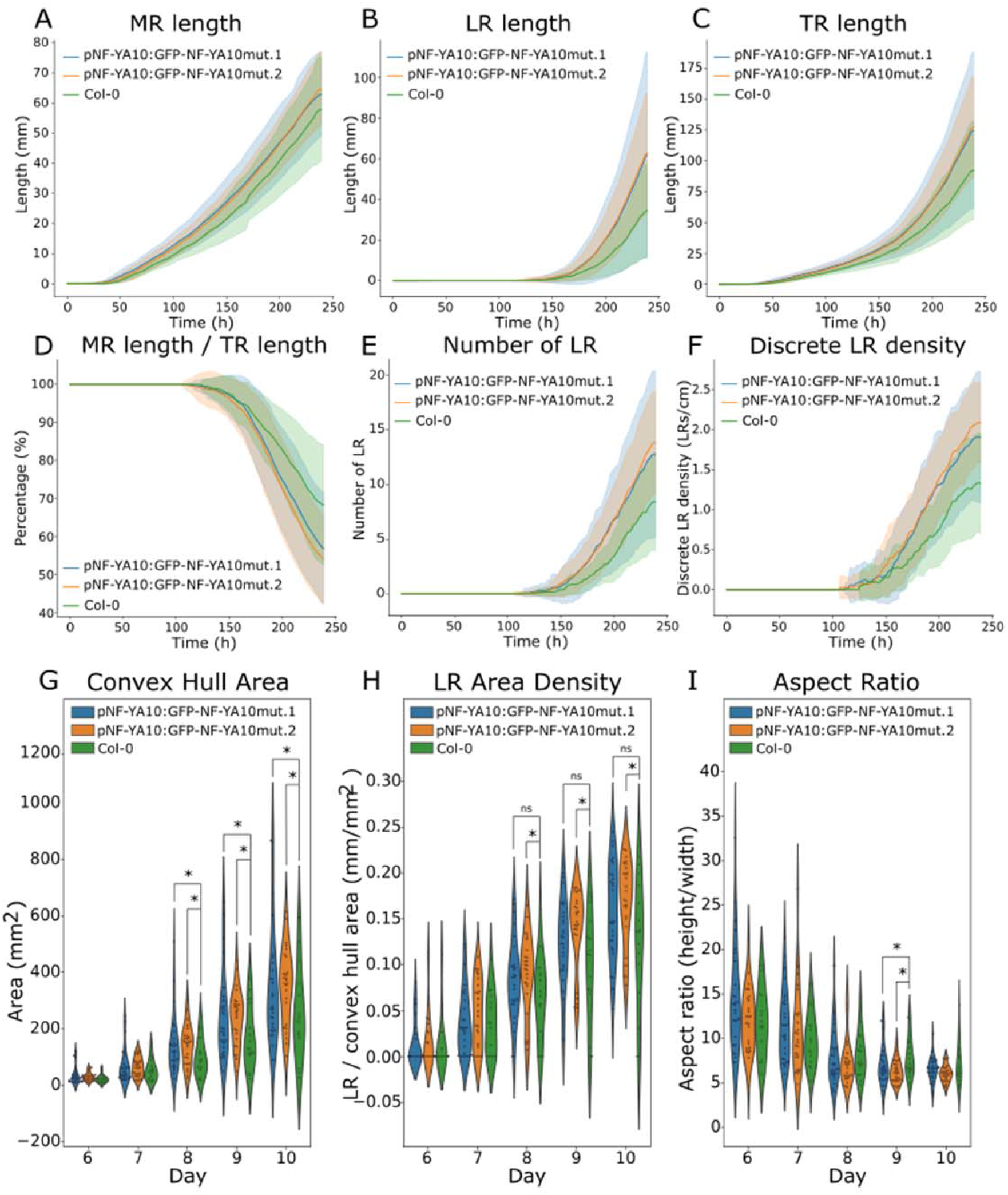
NF-YA10 up-regulation affects root growth and the resulting root system architecture. ChronoRoot measurements of NF-YA10 miRres and Col-0 WT plants: main root length (A), lateral root length (B), total root length (C), ratio main root length/total root length (D), number of lateral roots (E), lateral root density (F). Convex hull area (G), lateral root area density (H) and ratio of root system height and width (I) of NF-YA10 miRres lines and Col-0 at different ages of the plant. A-F: solid line represent the average value and the bands represent the standard deviation.

This characteristic feature led us to wonder if the angle of the LRs was affected in NF-YA10 miRres plants. To address this issue, two novel parameters were incorporated into ChronoRoot by leveraging information from both the segmentation and graph structures generated by the deep learning-based model. These parameters allowed for tracking the evolution of the base-tip angle (from the base to the tip of each LR) and the emergence angles of LRs over time (**Figure 3A-C, Supplementary Figure 2**), thereby expanding the original ChronoRoot applications. Considering the first LR emerged per plant, NF-YA10 miRres seedlings present a higher base-tip angle than the WT during the emergence of LR, reaching a 20°-difference after three days (**Figure 3D**). Accordingly, the emergence angle of LR was also affected, as NF-YA10 miRres plants showed a trend of greater emergence angles than the WT starting from the emergence of the first lateral roots (day 6), which was statistically significant for both independent lines from day 9 on (**Figure 3E**). The potential role of NF-YA10 in gravitropism was also assessed in the MR growth during a bending assay (i.e. plant plates turned 90° and MR growing angles measured afterwards). Interestingly, one of the two NF-YA10 miRres lines showed a significantly delayed response compared to WT suggesting that the role of this TF in LR development may also be relevant in the response of the MR to gravitropic stimuli (**Supplementary Figure 3**).

**Figure 3.**
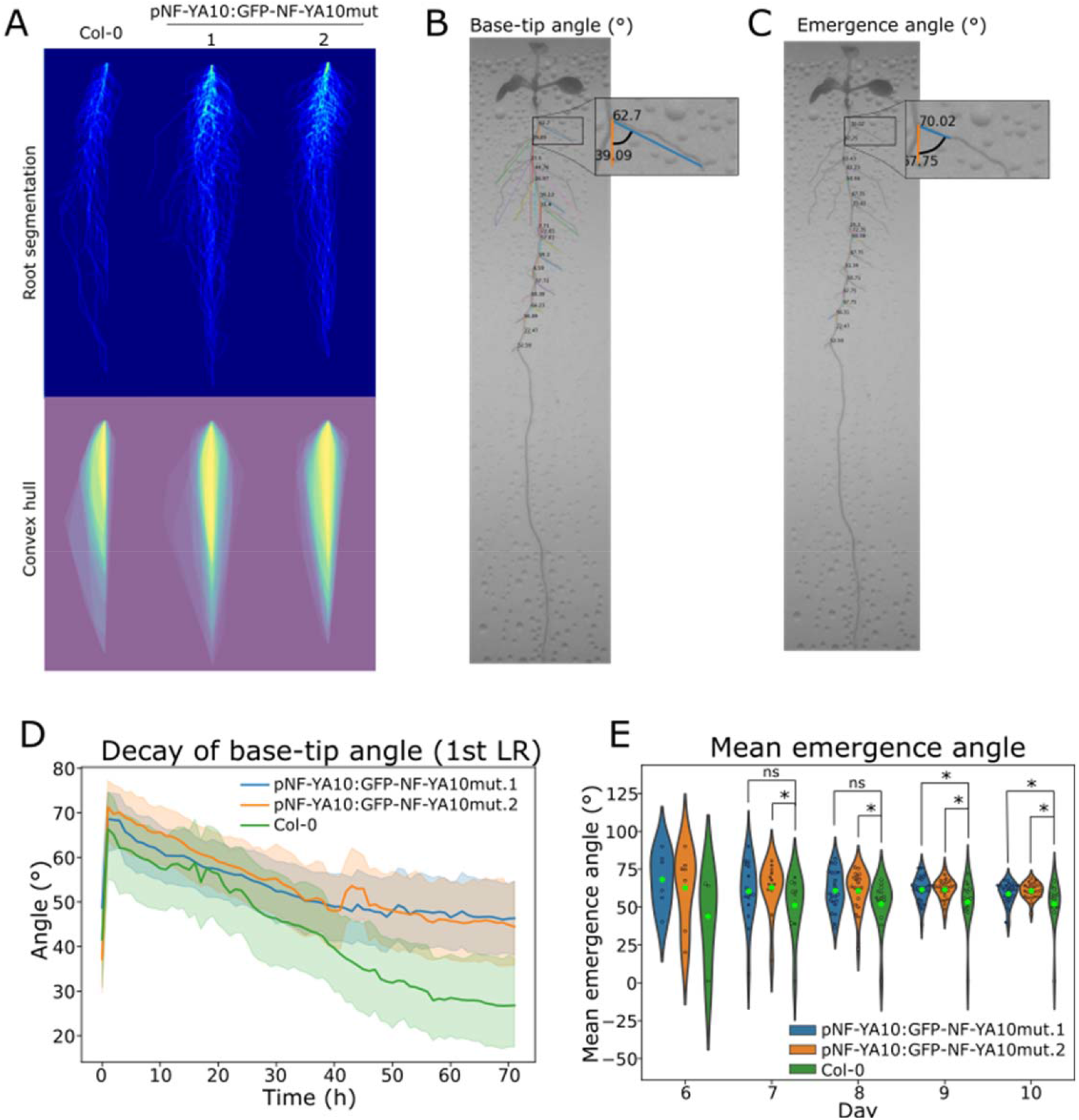
Two new features of ChronoRoot allowed us to uncover NF-YA10 as a regulator of lateral root gravitropism by leveraging information from both the segmentation and graph structures generated by the system. (A) Superposition of all NF-YA10 miRres and Col-0 root profiles in mock conditions. Representative plant of NF-YA10 miRres and zoom on two novel ChronoRoot measurements: (B) base-tip angle and (C) emergence angle respectively. (D) Dynamics of tip decay of the first lateral root to emerge along the time of NF-YA10 miRres plants and Col-0. (E) Mean emergence angle of NF-YA10 miRres and Col-0 roots at different ages of the plant.

To decipher the molecular mechanism behind this phenotype, we looked for potential NF-YA10 targets involved in lateral organ gravitropism described in the literature, which bear CCAAT-boxes in their promoter sequences. To this end, we first crossed the list of differentially expressed genes (DEGs) in NF-YA10-inducible overexpressor (Leyva-González et al., 2012) with TAIR Gene Ontology lists for genes related positively or negatively to gravitropism (**Supplementary Table 2**). Among them, we identified the genes *LAZY2* and *GLV9*, and *PIF8* respectively (**Figure 4A**). In contrast to *GLV9* and *PIF8, LAZY2* was transcriptionally deregulated in roots of NF-YA10 miRres plants (**Figure 4B**). Considering the behavior of *LAZY2* in NF-YA10-deregulated plants, we further investigated the transcriptional levels of members of the LAZY protein-encoding gene family, which have been linked to redundant activity (Hollander *et al*., 2020). Particularly, *TAC1* (*TILLER ANGLE CONTROL 1*) and *LAZY1/2/3* are known to be essential for accurate gravitropic auxin-driven sensing of roots and shoots. Interestingly, it was shown that LRs in the *lazy1/lazy2/lazy3* triple mutant display a disturbed gravitropic response whereas *tac1* simple mutant has an increased gravitropic phenotype (Hollander *et al*., 2020). Thus, we assessed their level of expression in NF-YA10 miRres roots. *LAZY1, LAZY2* and *LAZY3* were all repressed compared to WT while *TAC1* was induced (**Figure 4C**), in agreement with the mutant gravitropic phenotypes observed. In order to determine if NF-YA10 directly regulates these genes, we performed ChIP-qPCR targeting TSS-proximal CCAAT boxes present in their promoter regions, compared to ChIP enrichment in their respective gene bodies taken as the negative control (probes distribution indicated in **Figure 4D**). In basal conditions, *LAZY1, LAZY2* and *TAC1* emerged as direct targets of NF-YA10 (**Figure 4E**). *LAZY3* appears only as a potential direct target, considering that the negative control within the gene body is close to the CCAAT box assessed. Altogether, our results suggest that regulation of NF-YA10 expression is critical for the control of the root system architecture, notably determining the final volumetric root distribution by modulating root growth, LR development and their gravitropic response. NF-YA10 recognizes the promoters and regulates a subset of root developmental genes, including the antagonistic *TAC1* and *LAZY* genes, known to be involved in LR gravitropic signaling suggesting it maybe a new regulatory circuit controlling the long-term surface of the root system (**Figure 4F**).

**Figure 4.**
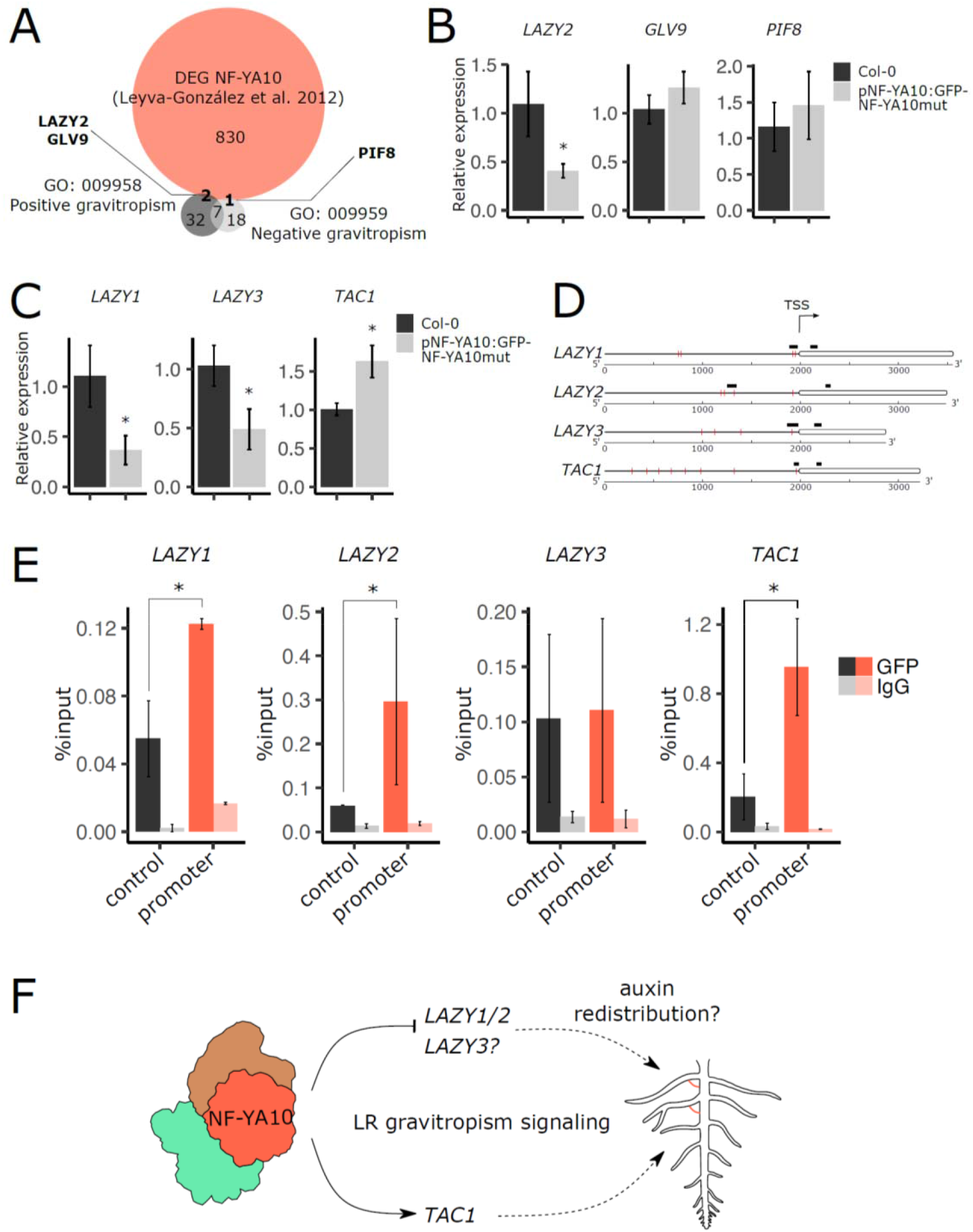
NF-YA10 binds to *TAC1* and *LAZYs* promoters, resulting in antagonistic regulation of these genes. (A) Venn diagram of DEGs from Leyva-González et al., 2012 and genes comprised in Gene Ontology terms of positive and negative gravitropism. (B) Location of CCAAT boxes are indicated in red lines and the respective DNA regions assessed in ChIP-qPCR in black for each of the genes *TAC1, LAZ1, LAZY2* and *LAZY3*. (C) Relative expression of *TAC1* and *LAZYs* in NF-YA10 miRres and Col-0 roots of 8-day-old seedlings. (D) Chromatin immunoprecipitation (ChIP)-qPCR analysis of NF-YA10 binding at *TAC1* and *LAZYs* gene and promoter regions in 8-day-old NF-YA10 miRres and Col-0 seedlings. (E) Proposed model for the NF-YA10-dependent lateral root gravitropic response. In (B), (C) and (E), values correspond to the mean and error bars to the standard deviation of three biological replicates. The asterisks indicate a p-value < 0.05 (Mann-Whitney test).

## DISCUSSION

In animals, the NF-Y complex is described as a pioneer TF (Dolfini et al., 2016), able to recruit chromatin remodelers and downstream TFs to modulate the activity of target genes. In Arabidopsis, NF-YA, NF-B and NF-YC are encoded by respectively 10, 13 and 13 genes boosting the diversification of molecular and physiological roles of this TF family, based on the wide range of combinations likely occurring in different cell types from the organism (Laloum et al., 2013). Among them, NF-YA2 and NF-YA10 emerged from a recent Brassicaceae-specific duplication inside a clade with low divergence degree, thus exhibiting few changes in the protein sequence. Both NF-YA genes are targets of the specific isoform of miR169-defg, and plants resistant to the downregulation driven by this miRNA showed similar LR phenotype (Sorin *et al*. 2014; and **Figure 2**). For a better characterization of NF-YA10 miRres plants, two novel root parameters were determined by ChronoRoot through the development of ad hoc deep segmentation networks: base-tip angle and emergence angle of LRs. These measures were respectively 20° and 8° higher in NF-YA10 miRres lateral roots than in the WT, causing, together with higher LR density, a significant increase of the area occupied by the root system in two dimensions over the agar plate. Strikingly, Arabidopsis WT roots exhibit a similar phenotype in response to phosphate starvation (Bai *et al*., 2013) and it was previously shown that the promoter of *NF-YA10* is induced under these conditions, whereas miR169 was repressed (Leyva-González *et al*., 2012). Our observations suggest that NF-YA10 may participate in the control of LR gravitropism in response to low phosphate concentrations. Considering that root-specific targets of NF-YA10 are yet unknown, we searched for potential targets that could be linked to the regulation of LR gravitropism, among which we identified *TAC1* and *LAZYs* genes, described as components of the signaling pathway controlling gravitropism in shoots and roots (Kawamoto and Morita, 2022).

All four genes assessed turned out to be up- or down-regulated by NF-YA10 in NF-YA10 miRres plants. In contrast to *LAZY1, LAZY2* and *TAC1* determined by ChIP-qPCR, *LAZY3* was not significantly enriched by NF-YA10 compared to the nearby negative control. Further research will be required to determine how this regulation may take place under phosphate starvation. Notably, lugol-staining of LRs revealed no observable changes in amyloplast distribution in NF-YA10 miRres roots, suggesting that NF-YA10 may operate downstream the known gravity sensing organelle (**Supplementary Figure 4)**. Lower levels of auxin were detected in NF-YA10 OE leaves, and a microarray transcriptomic approach of the same plants served to determine that the expression of a significant subset of auxin-related genes depends on NF-YA10 (Zhao *et al*., 2017), further suggesting that this TF may play an important role in auxin homeostasis, likely explaining the phenotypes observed in root growth, development and LR gravitropism. Furthermore, the use of the synthetic reporter DR5 allowed demonstrating that auxin signaling was increased in triple *lazy2/lazy3/lazy4* mutant roots (Yoshihara & Spalding, 2017; Ge & Chen, 2019), further linking the impact of the NF-YA10-LAZYs hub on auxin-driven root development. In addition, the ortholog of Arabidopsis *LAZY1* in maize, ZmLAZY1, directly interacts with the early auxin response factor ZmIAA17 in the nucleus of maize cells (Dong et al., 2013). In Arabidopsis, the dwarf late-senescent phenotype of NF-YA10 overexpressing plants under the control of a 35S promoter, was previously proposed to be caused by a reduction of plant growth directed by transcriptional regulation of cell wall and sucrose pathways actors (Leyva-González *et al*., 2012). In contrast to 35S-mediated overexpression, a miR-resistant version induces a slight up-regulation and in the same cells expressing the NF-YA10 TF. Therefore, we propose that NF-YA10 contributes to orchestrate LR development in addition to auxin and cell expansion related genes, by fine-tuning gravitropism signaling through regulation of *LAZY* genes expression. Indeed, NF-YA10 deregulation results in a larger root area. It was shown that enhanced root surface area positively correlated with plant weight, opening new perspectives about the potential of NF-Y TFs for the improvement of relevant traits for hydroponics culture and more globally for agriculture (Yang *et al*., 2021). Therefore, further research about how LRs explore the substrate under the control of transcriptional regulators could help to improve crop growth based on nutrients uptake, including phosphates.

## MATERIAL AND METHODS

### Plant lines generated and used for this study

All plants used in this study are in Columbia-0 background. pNF-YA10:GFP-NF-YA10miRres lines were obtained using the GreenGate vectors (Lampropoulos *et al*., 2013), using 2000-pb region upstream of the start codon of *NF-YA10* amplified from genomic DNA and CDS of NF-YA10 without miR cleavage site amplified from cDNA. Arabidopsis plants were transformed by floral dip (Clough and Bent 1998) using *Agrobacterium tumefaciens* C58. Homozygous mutants of miR169 resistant line of NF-YA10 were identified by PCR (see primers in **Supplementary Table 3**).

### Growth conditions and phenotypic analyses

For phenotype analyses, plants were grown vertically on plates placed in a growing chamber in long day conditions (16 h in light 130uE; 8 h in dark; 23°C). Plants were grown on solid half-strength MS medium (MS/2) supplemented with 0.8g/L agar (Sigma-Aldrich, A1296 #BCBL6182V) and 1% sucrose, buffered at pH 5,6. Temporal phenotyping was performed using ChronoRoot (Gaggion et al., 2021). In the case of gravitropic assays, plates were rotated at 90° when reaching 7 DAS and photographed after 24 hours. Images where then analyzed with ImageJ to measure the root tip angle of each plant (n=15) formed after reorientation. For the observation of amyloplasts, 8-day-old plants (n=15) were dipped for 8 minutes in Lugol staining solution (0.1% I, 1%KI, Sigma-Aldrich) and observed in an Eclipse E200 Microscope (Nikon) equipped with a Nikon D5300 camera.

### Confocal laser scanning and Fluorescence microscopy (CLSM)

For CLSM, roots of stable two independent lines of NF-YA10 miRres plants were imaged with a Leica TCS SP8 confocal laser scanning microscope. For GFP signal imaging, samples were excited at 488 nm and the detection was set at 493–530nm, and the transmitted light was also imaged. All the images were captured using a 20x or 63x lens. Image processing was performed using Fiji software (Schindelin et al. 2012).

### Sequence alignment and phylogenetic tree analysis

Protein sequences corresponding to plant NF-YA family members were identified using the BLASTP tool (Boratyn et al., 2013) and downloaded from Phytozome 13 (https://phytozome-next.jgi.doe.gov/) (Goodstein et al., 2012). Proteins from other organisms were obtained from the NCBI database. For the tree in **Supplementary Figure 1**, protein sequences of plant species were selected by choosing members from the main phylogenetic clades, giving a total of 81 protein sequences (**Supplementary Table 1**). Three no-plant species sequences were using for tree rooting. For the tree in **Figure 1A**, several protein sequences from Brassicaceae and Brassicales-Malvales species were added to improve resolution in this group of organisms, resulting in a total of 137 protein sequences (**Supplementary Table 1**). The alignments were made using MUSCLE default parameters (Edgar, 2004). The phylogenetic trees were built using the Seaview 4.5.0 software and the PhyML-aLRT-SH-LIKE algorithm (Gouy et al., 2010) with maximum likelihood tree reconstruction. A model of the amino acid substitution matrix was chosen through the Datamonkey bioinformatic server (www.datamonkey.org; Delpor *et al*., 2010), which selected the VT model. The resulting trees were represented using iTOL (http://itol.embl.de/itol.cgi; (Letunic & Bork, 2016)) showing branches with bootstrap values higher than 65%.

### Quantification of transcript levels by RT-qPCR

Total RNA was extracted from roots using TRI Reagent (Sigma-Aldrich) and treated with DNaseI (NEB) as indicated by the manufacturers. Reverse transcription was performed using 1μg total RNA and the M-MLV Reverse Transcriptase (Promega). qPCR was performed on a StepOne™ Real-Time PCR System (Thermo Fisher) with Sso Advanced Universal mix (BioRad) in standard protocol (40 cycles, 60°C annealing). Primers used in this study are listed in **Supplementary Table 3**. Data were analyzed using the ΔΔCt method using ACTIN (AT3G18780) for gene normalization (Czechowski et al., 2005).

### Chromatin immunoprecipitation

Three biological replicates of 8 DAS whole seedlings grown in control condition were collected. ChIP was performed using anti-GFP (Abcam ab290) and anti-immunoglobulin G (IgG) anti-IgG (Abcam ab6702), as described in Ariel et al. 2020, starting from 5 g of seedlings crosslinked in 1% (v/v) formaldehyde. Chromatin was sonicated in a water bath Bioruptor Plus (Diagenode; 10 cycles of 30 s ON and 30 s OFF pulses at high intensity). Antibody-coated Protein A Dynabeads (Invitrogen) were incubated 12 h at 4°C with the samples. Immunoprecipitated chromatin was reverse crosslinked with 20 mg of Proteinase K (Thermo Fisher, #EO0491) overnight at 65°C. Finally, DNA was recovered using phenol/chloroform/isoamilic acid (25:24:1, Sigma) followed by ethanol precipitation. For input samples, 10% of sonicated chromatin were collected for each sample before the immunoprecipitation and reverse crosslinked and extracted as the immunoprecipitated samples. Results are expressed as enrichments, corresponding to GFP or IgG percent of input, measured by qPCR (primers used are listed in **Supplementary Table 3**).

### Construction of Angle Parameters based on ChronoRoot

To construct the new angle parameters, specifically the base-tip angle (**Figure 3B**) and the emergence angle of LR (**Figure 3C**), we first proceeded on extracting the LRs from the labeled skeleton of each plant. Then, to preserve the labels and information pertaining to the order in which the LRs began to grow, we matched the extracted LRs across time-steps. We performed the matching from one time-step to the previous one by using the pixel position where the root initiated. With the position, skeleton, and label of each plant root established, we proceeded to measure the LR base-tip angle. This angle was calculated for each LR by examining the right triangle formed by the base position, the tip, and the Y-axis. For the second angle parameter, i.e. the emergence angle, instead of measuring it from the base to the tip, we established a fixed distance in millimeters (2 mm in our study, though adjustable as a parameter) to construct the right triangle (see **Figure 3C**). We traversed 2 mm along the skeleton from the base and measured the angle between the Y-axis and this point to calculate it. Source code for ChronoRoot is publicly available on GitHub (https://github.com/ngaggion/ChronoRoot, Gaggion et al., 2021).

## FIGURE LEGENDS

**Figure 1. NF-YA10 diverged from a recent duplication with NF-YA2 within Brassicaceae and is expressed in nuclei of primary and lateral root vasculature cells**

(A) Phylogenetic tree of NF-YAs with extended number of Malvidae and Brassicaceae, where AtNF-YA10 is represented in red. (B) Localization of NF-YA10 fused to GFP in roots of pNF-YA10:GFP-NF-YAmiRres.1 8-day-old plants.

**Figure 2. NF-YA10 up-regulation affects root growth and the resulting root system architecture**

ChronoRoot measurements of NF-YA10 miRres and Col-0 WT plants: main root length (A), lateral root length (B), total root length (C), ratio main root length/total root length (D), number of lateral roots (E), lateral root density (F). Convex hull area (G), lateral root area density (H) and ratio of root system height and width (I) of NF-YA10 miRres lines and Col-0 at different ages of the plant. A-F: solid lines and corresponding bands represent the average value and the standard deviation respectively. G-I: The asterisks indicate a p-value < 0.05 (Mann-Whitney test).

**Figure 3. Two new features of ChronoRoot allowed us to uncover NF-YA10 as a regulator of lateral root gravitropism by leveraging information from both the segmentation and graph structures generated by the system**

(A) Superposition of all NF-YA10 miRres and Col-0 root profiles in mock conditions. Representative plant of NF-YA10 miRres and zoom on two novel ChronoRoot measurements: (B) base-tip angle and (C) emergence angle respectively. (D) Dynamics of tip decay of the first lateral root to emerge along the time of NF-YA10 miRres plants and Col-0. Solid lines and corresponding bands represent the average value and the standard deviation respectively. (E) Mean emergence angle of NF-YA10 miRres and Col-0 roots at different ages of the plant. The asterisks indicate a p-value < 0.05 (Mann-Whitney test).

**Figure 4. NF-YA10 binds to *TAC1* and *LAZYs* promoters, resulting in antagonistic regulation of these genes**

(A) Venn diagram of DEGs from Leyva-González et al., 2012 and genes comprised in Gene Ontology terms of positive and negative gravitropism. (B) Location of CCAAT boxes and the respective DNA probes used in ChIP-qPCR for the study of the genes *TAC1, LAZ1, LAZY2* and *LAZY3*. (C) Relative expression of *TAC1* and *LAZYs* in NF-YA10 miRres and Col-0 roots of 8-day-old seedlings. (D) Chromatin immunoprecipitation (ChIP)-qPCR analysis of NF-YA10 binding at *TAC1* and *LAZYs* gene and promoter regions in 8-day-old NF-YA10 miRres and Col-0 seedlings. (E) Proposed model for the NF-YA10-dependent lateral root gravitropic response. In (B), (C) and (E), values correspond to the mean and error bars to the standard deviation of three biological replicates. The asterisks indicate a p-value < 0.05 (Mann-Whitney test).

## DISCLOSURES

### Ethics approval and consent to participate

Not applicable for this study.

## SUPPLEMENTARY FIGURES

**Figure S1**. Phylogenetic tree of NF-YAs in plant and non-plant organisms, where AtNF-YA10 is represented in red.

**Figure S2**. Representative plants of NF-YA10 miRres (line 2) and WT with corresponding measures of base-tip and emergence angles.

**Figure S3**. (A) Representative photography of 8-day-old NF-YA10 miRres and Col-0 seedlings and (B) mean angle of corresponding primary roots 48 hours after gravistimulation.

**Figure S4**. Representative photography of (A) primary and (B) lateral root apexes of 8-day-old NF-YA10 miRres and Col-0 seedlings showing amyloplasts stained with lugol.

## SUPPLEMENTARY TABLES

**Table S1**. Table of species and sequences used for phylogenetic analyses.

**Table S2**. Lists of differentially expressed genes (DEGs) in *NF-YA10*-inducible overexpressor (Leyva-González et al., 2012) and genes assigned to TAIR Gene Ontology terms related positive and negative gravitropism.

**Table S3**. Table of primers used in this study.

